# Methods for Processing and Analyzing Images of Vascularized Micro-Organ and Tumor Systems

**DOI:** 10.1101/2025.03.12.642741

**Authors:** Stephanie J. Hachey, Christopher J. Hatch, Daniela Gaebler, Alexander G. Forsythe, Makena L. Ewald, Alexander L. Chopra, Jennifer S. Fang, Christopher C.W. Hughes

## Abstract

Our group has developed and validated an advanced microfluidic platform to improve preclinical modeling of healthy and disease states, enabling extended culture and detailed analysis of tissue-engineered miniaturized organ constructs, or “organs-on-chips.” Within this system, diverse cell types self-organize into perfused microvascular networks under dynamic flow within tissue chambers, effectively mimicking the structure and function of native tissues. This setup facilitates physiological intravascular delivery of nutrients, immune cells, and therapeutic agents, and creates a realistic microenvironment to study cellular interactions and tissue responses. Known as the vascularized micro-organ (VMO), this adaptable platform can be customized to represent various organ systems or tumors, forming a vascularized micro-tumor (VMT) for cancer studies. The VMO/VMT system closely simulates in vivo nutrient exchange and drug delivery within a 3D microenvironment, establishing a high-fidelity model for drug screening and mechanistic studies in vascular biology, cancer, and organ-specific pathologies. Furthermore, the optical transparency of the device supports high-resolution, real-time imaging of fluorescently labeled cells and molecules within the tissue construct, providing key insights into drug responses, cell interactions, and dynamic processes such as epithelial-mesenchymal transition. To manage the extensive imaging data generated, we created standardized, high-throughput workflows for image analysis. This manuscript presents our image processing and analysis pipeline, utilizing a suite of tools in Fiji/ImageJ to streamline data extraction from the VMO/VMT model, substantially reducing manual processing time. Additionally, we demonstrate how these tools can be adapted for analyzing imaging data from traditional *in vitro* models and microphysiological systems developed by other researchers.

## 1 INTRODUCTION

Preclinical organ-on-a-chip models that closely replicate human physiology and pathology – and especially the blood vasculature – are indispensable to advance disease research, drug discovery, and personalized medicine (Hachey and Hughes (2018); Low and Tagle (2018); Low et al. (2021); Ewald et al. (2021); Ingber (2022); Martier et al. (2024); Gaebler et al. (2024a,b)). To address the limitations of existing models that fail to recapitulate a vascularized tissue niche, we developed the vascularized micro-organ (VMO) platform, an advanced organ-on-a-chip system that supports long-term studies of tissue-engineered, miniaturized organ constructs with associated microvasculature. This dynamic microfluidic platform allows for the co-culture of multiple cell types in a controlled flow environment, enabling the self-assembly of perfused microvascular networks within 3D tissue chambers. The physiological relevance of the platform is further enhanced by its ability to deliver nutrients and therapeutic agents through functional vascular networks, creating a powerful tool for studying vascular biology, disease progression, and therapeutic responses in vitro (Phan et al. (2017a); Urban et al. (2018); Bender et al. (2024); Hatch et al. (2024)).

One application of the VMO platform is the vascularized micro-tumor (VMT) model, which integrates tumor cells and stromal components into a 3D extracellular matrix within the tissue chambers. Gravity-driven fluid flow facilitates the rapid formation of living, perfused microvascular networks that support tumor growth and drug delivery, closely mimicking the complexity of in vivo tumor biology (Sobrino et al. (2016); Phan et al. (2017b); Hachey et al. (2021, 2022, 2023, 2024)). The VMO/VMT system uniquely recreates the stromal-vascular interactions critical to understanding disease mechanisms and evaluating therapeutic strategies, overcoming many of the limitations of conventional drug-screening models.

Given the intricate spatial and temporal data generated from imaging experiments with the VMO/VMT platform, efficient and reproducible data analysis workflows are essential. To address this need, we developed Hughes Lab Tools, a suite of custom-designed image-processing algorithms implemented in ImageJ/Fiji, an open-source Java-based image processing program developed by the National Institutes of Health (Schindelin et al. (2012)). ImageJ’s open architecture enables extensibility through Java plugins, recordable macros, and scripts written in various programming languages. Using these capabilities, Hughes Lab Tools incorporates user-friendly Jython scripts and ImageJ macros to automate and standardize image processing for VMO/VMT tissue constructs.

This set of tools enables high-throughput data extraction and automates critical tasks such as quantifying tumor growth, vascular remodeling, and flow dynamics. Hughes Lab Tools supports both fully automatic and semi-automatic workflows, allowing operators to verify intermediate results when necessary. Images can be processed from single directories or nested folder structures, and users can execute individual functions or run multiple tasks in series, vastly improving throughput over manual methods. Moreover, these tools are easily modifiable, offering flexibility to address a broad range of experimental questions beyond the VMO/VMT platform.

Here, we present the design and application of the Hughes Lab Tools suite on data generated from the VMO/VMT platform, demonstrating how it streamlines data extraction and analysis while maintaining accuracy and reproducibility. Furthermore, we highlight the broad applicability of Hughes Lab Tools for image-based analysis in other preclinical model systems, including organ-on-a-chip technologies and microphysiological platforms, making them a valuable resource for diverse areas of biomedical research.

## 2 MATERIALS AND EQUIPMENT

1. Computation
  - ImageJ/Fiji software
  - Autodesk software
  - COMSOL Multiphysics software
2. Equipment
  - Biotek Lionheart automated fluorescent microscope
  - Leica SP8 confocal microscope
3. Materials
  - Cell culture medium
  - Cell culture reagents
  - Endothelial cells
  - Stromal cells
  - Cancer cells (*optional*)

## 3 METHODS

### 3.1 Script Development

The development of the tool suite was managed using Git version control (Community (2025)), with the complete version history and code accessible on GitHub (https://github.com/aforsythe/HughesLabTools). Script development was conducted within the ImageJ and Fiji distributions (Schindelin et al. (2012)), which provide a suite of tools for creating macros and scripts (Schneider et al. (2012)). These distributions include a Macro Recorder to assist in identifying command sequences for user interface operations. An integrated development environment (IDE) was employed for its advanced coding and debugging capabilities. Specifically, IntelliJ IDEA Community Edition (JetBrains) was utilized. This Java Virtual Machine (JVM)-based IDE (Lindholm et al. (2014)) supports multiple programming languages, including Java and Python. Python served as the primary language for script development due to its compatibility with ImageJ and Fiji.

### 3.2 Cell Culture

Human endothelial colony-forming cell-derived endothelial cells (ECFC-EC) were isolated from cord blood following an IRB-approved protocol. After selecting for the CD31+ population, ECFC-ECs were expanded in gelatin-coated flasks using EGM2 medium (Lonza) and used between passages 4 and 8. Normal human lung fibroblasts (NHLFs) were procured from Lonza and utilized between passages 6 and 10. The human non-small cell lung cancer line H1792 was obtained from ATCC. ECFC-ECs and cancer cells were transduced with lentiviruses expressing mCherry (LeGO-C2, Addgene plasmid #27339), green fluorescent protein (GFP; LeGO-V2, plasmid #27340), or azurite (pLV-azurite, plasmid #36086). NHLFs and cancer cells were cultured in DMEM (Corning) or RPMI-1640 supplemented with 10% FBS (Gemini Bio). All cells were maintained at 37 °C and 5% CO_2_.

### 3.3 Tumor Spheroid Generation & Seeding

Tumor spheroid formation was performed using AggreWell™ plates (StemCell Technologies) to promote the aggregation of H1792 lung cancer cells. Briefly, H1792 cells were seeded at a density of 2 *×* 10^5^ cells per well in 500 μL of complete DMEM (Dulbeccos Modified Eagle Medium) supplemented with 10% fetal bovine serum (FBS). The cells were allowed to aggregate in the AggreWell™ plates for 48 h at 37°C and 5% CO_2_ incubator. After 48 h, the spheroids were carefully harvested for downstream applications.

### 3.4 3D Spheroid Culture and Drug Treatment

H1792 cancer cells were suspended in a 5 mg*/*mL fibrinogen solution at a density of 1 × 10^6^ cells/mL. Fifty microliters of the cell suspension were added to each well of a 96-well plate containing 5 U of thrombin. The mixture was allowed to polymerize at 37 °C for 15 min. Following polymerization, 100 μL of EGM2 medium was added to each well. Drug treatments were initiated 6 h post-seeding and were maintained for 48 h. After 48 h, the drug-containing medium was removed and replaced with fresh EGM2 medium. Fluorescent micrographs of the spheroids were captured every 48 h to monitor changes in spheroid morphology and evaluate drug responses.

### 3.5 Microfluidic Device Fabrication

Device fabrication followed previously described methods (Sobrino et al. (2016); Phan et al. (2017b); Hachey et al. (2023)). Briefly, polydimethylsiloxane (PDMS) was prepared by mixing Sylgard 184 elastomer base with curing agent (10:1 ratio, Dow Corning), degassing the mixture, and casting it into a polyurethane master mold derived from a lithographically patterned silicon wafer. PDMS cast into the mold was cured at 70 °C for 4 hours, after which inlets and outlets were punched and the platform was assembled in two steps. First, the PDMS layer was bonded to the base of a 96-well plate using chemical glue and oxygen plasma treatment. Second, a 150 μm-thin transparent membrane was attached to the bottom of the PDMS layer after additional plasma treatment. Assembled devices were cured overnight at 70 °C, sterilized with UV light for 30 minutes, and stored until cell loading.

### 3.6 Establishment of vascularized micro-organ (VMO) and vascularized micro-tumor (VMT) Models

Establishment of the VMO and VMT models was performed according to published methods (Hachey et al. (2023)). Briefly, to establish the VMO, endothelial colony-forming cell-derived endothelial cells (ECFC-EC)s and normal human lung fibroblasts (NHLF)s were resuspended in a 10 mg*/*mL fibrinogen solution at a density of 7 × 10^6^ cells/mL and 3.5 × 10^6^ cells/mL, respectively. For the VMT, lung cancer cells were added to this fibrinogen mixture at 2 × 10^5^ cells/mL. Fibrinogen solution was prepared by dissolving 70% clottable bovine fibrinogen (Sigma-Aldrich) in EBM2 basal medium (Lonza) to a final concentration of 5 mg/mL. The cell-matrix suspension was mixed with thrombin (50 U/mL, Sigma-Aldrich) to achieve a final concentration of 3 U/mL and immediately introduced into microtissue chambers. Polymerization occurred in a 37 °C incubator for 15 minutes. Laminin (1 mg/mL, LifeTechnologies) was introduced into the microfluidic channels to support vessel anastomosis and incubated for 15 minutes before replacing with culture medium (EGM-2). Media reservoirs were filled with EGM2 to establish a hydrostatic pressure head. Medium changes were conducted every other day, while levels were adjusted daily to maintain interstitial flow.

### 3.7 Drug Treatment in the VMT

After 4-5 days of culturing, a perfused vascular network was established within each VMT, and the culture medium was replaced with drug-containing medium at the desired concentrations. Drug delivery to the tumor was achieved through the vascular bed via gravity-driven flow. Paclitaxel (a microtubule stabilizer) was purchased from SelleckChem. For H1792 VMTs, experimental groups were randomly assigned to one of three conditions: control (vehicle only), 200 nM paclitaxel, or 400 nM paclitaxel. Oregon green 488-conjugated paclitaxel was purchased from Invitrogen. The medium was replaced after 48 hours. Fluorescent micrographs of VMTs were taken every 48 hours for six days post-treatment, and tumor growth was quantified.

### 3.8 Fluorescence Imaging and Perfusion

Fluorescence imaging was conducted using a Biotek Lionheart fluorescent inverted microscope with automated acquisition and a standard 10X air objective. Vessel perfusion and permeability were evaluated by adding 25 μ*g/*mL FITC- or rhodamine-conjugated 70 kDa dextran to a medium inlet. Time-lapse image sequences were acquired once the fluorescent dextran reached the vascular network. Images were taken once every minute for 20 minutes. VMOs were treated with 100 ng/mL VEGF_165_ for 24 hours prior to perfusion to assess changes in permeability. Both VMOs and VMTs were perfused on day 5–6 of culture, when a complete vascular network had formed. Confocal imaging was performed on a Leica TCS SP8 confocal microscope using a standard 10X air objective or 20X multi-immersion objective with digital zoom setting.

### 3.9 Image Segmentation Using WEKA in Fiji

The Trainable Weka Segmentation plugin in Fiji/ImageJ was used for image segmentation. The plugin was installed via Fijis Update feature, and segmentation was performed by opening the image and launching Trainable Weka Segmentation. A Feature Set was selected, and the Brush Tool was used to manually annotate different regions. The Train Classifier function refined segmentation based on user-labeled samples. The trained model was saved and applied to new images via File – Load Classifier and – Apply Classifier. The final segmentation was generated using Create Probability Map, refined through Thresholding and Morphological Operations (Fill Holes, Watershed). Processed images were saved for analysis.

### 3.10 Finite Element Simulations

Finite element modeling of interstitial flow within vascular networks was performed in COMSOL Multiphysics 5.2a (COMSOL AB (2023)). Vessel images processed in ImageJ were converted into.dxf files using the Hughes Lab Tools with custom code based on MATLAB DXFLib (Kwiatek (2025)) and refined in AutoCAD for integration into a 2D flow model. Culture media flow was modeled as water, with fibrin gel properties set to a porosity of 0.99 and a permeability of 1.5 × 10^*−*13^ m^2^ (Helm et al. (2005); Hsu et al. (2013)). Pressure gradients were applied based on gravity-driven flow parameters.

### 3.11 Image Analysis

Image processing and analysis were conducted using the Hughes Lab Tools script suite. This versatile suite facilitated the evaluation of area, circularity, and roundness for each tumor image, providing critical metrics for assessing tumor growth. All measurements were normalized to baseline levels. Tumor growth in the VMTs was assessed by analyzing total fluorescence intensity (mean gray value), circularity/roundness, and area within the color channel corresponding to tumor cells. This analysis accounted for both tumor area and depth, with thicker regions exhibiting higher brightness due to increased fluorescence intensity. Similarly, tumor spheroid growth was monitored by tracking changes in fluorescence intensity, spheroid roundness, and tumor area over time. Vessel parameters including area, length, diameter, junctions, and endpoints were quantified using the Hughes Lab Tools suite. All measures were normalized to their baseline values to enable accurate longitudinal comparisons.

Vessel permeability was assessed by fluorescence changes in extravascular regions as analyzed by selecting multiple regions of interest (ROI) per image. Permeability coefficients were calculated as described in Eq. 1:

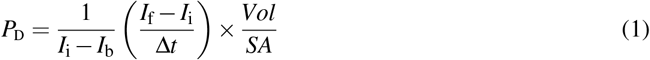

where *I*_i_, *I*_f_, and *I*_b_ represent the initial, final, and background intensities, respectively; Δ*t* is the time interval between images; *VOL* is the volume of the tissue; and *SA* is the surface area of the vessels (Campisi et al. (2018); Ahn et al. (2020); Hajal et al. (2022); Nahon et al. (2024)). *VOL*/*SA* was approximated as *d*/4, or vessel diameter/4. Perfusion images were further analyzed to calculate fluorescence intensity changes within perfused vessel regions, generating a composite score based on total perfusable vascular area. Image adjustments, where necessary, were applied consistently across the entire experiment to maintain uniformity.

### 3.12 Vessel Quantification

Vessel quantification was performed using a combination of built in ImageJ Commands and the AnalyzeSkeleton plugin. The input image was cleaned using a distance map before being skeletonized with the skeletonization ImageJ command. The number of junction points was determined with the AnalyzeSkeleton plugin. Junction points were filtered based on a distance threshold. To analyze each branch, the skeleton was broken at junction points and analyzed. The diameters were determined using a distance map. ImageJ’s particle analysis functions was used to determine the area and perimeter. Results are saved in CSV format, including a summary table and detailed skeleton values.

### 3.13 Statistical Analysis

Statistical analyses were conducted using GraphPad Prism (Version 10.4.1). Data are represented as mean ± standard deviation of at least three independent experiments. Comparison between experimental groups of equal variance was analyzed using an unpaired t-test and 95% confidence interval or two-way ANOVA followed by Dunnett’s test for multiple comparisons. Statistical significance was defined as *p <* 0.05.

### 3.14 Hughes Lab Tools User Guide

1. **Script Installation**
  a. The Hughes Lab Tools suite was validated in the Fiji ImageJ2 distribution (Schindelin et al. (2012)) on macOS 10.14.4. Users are encouraged to use Fiji due to its inclusion of plugins not typically available in the base ImageJ distribution. To install Fiji on macOS, use the Homebrew package manager (Howell et al. (2025)) with the command: brew cask install fiji
  b. To facilitate the installation of Hughes Lab Tools, a shell script is provided. Follow these steps:
    (1) Download the suite from GitHub and extract the compressed file.
    (2) Navigate to the directory containing the scripts and execute the installer script with:

~~~
./hugheslabtools_install.sh
~~~
    (3) Choose one of the following installation modes:
      - “Copy Hughes Lab Tools to Fiji (End-user Mode)”: Copies source files into the Fiji.app directory.
      - “SymLink Hughes Lab Tools to Fiji (Developer Mode)”: Creates symbolic links to the tools directory for easier development and testing.
      - “Remove Hughes Lab Tools”: Removes installed files.
  c. Once installed, a “Hughes Lab Tools” menu becomes accessible in Fiji’s menu bar. Users can create custom keyboard shortcuts using the “Add Shortcut…” option in the “Plugins > Shortcuts” menu.
2. **Workflow Overview**
  a. Select Functions to Execute
    - Start by selecting one or more image processing functions from the “Hughes Lab Tools” menu.
    - Click Next to proceed.
    - Please see Figure 1 for an outline of the graphical user interface.
  b. Specify Image Types in Sub-directories
    - Indicate the number of distinct image types present in the sub-directories (e.g., Vessel, Tumor).
    - Ensure all sub-directories to be processed contain the same number of image types.
  c. Image Classification Methods
    - One classification methods is currently available:
      1. Snake Pattern Image Classification: Assumes images follow a sequential pattern.
  d. Verify Image Classification (Optional)
    - Check the “Confirm Image Types” option to manually verify classifications during processing.
  e. Select Function-Specific Options
    - Adjust settings such as image coloring, output format, and processing verbosity via the options dialog.
  f. Choose Image Directories
    - Select the directory (and sub-directories) containing images to process. Images should be provided in TIF format.
  g. Process Images
    - Functions can run automatically or allow users to verify intermediate steps.
  h. Shortcut Scripts for Single Functions
    - Shortcut scripts are available for streamlined execution of single functions, bypassing the unified selection dialog.
3. **Tools and Modules**
  a. **Color and Merge Images** This module colors monochrome micrograph images (e.g., FITC- and mCherry-labeled cells) and optionally merges them into composite images. Steps:
    (1) Specify colors for each image type in the options dialog.
    (2) Colored images are saved in a “Colored” sub-directory as JPEG files.
    (3) If merging is selected, composite images are saved in a “Merged” sub-directory as “composite_#” files.
  b. **Crop Images** This module allows for cropping of images. Steps:
    (1) Select either the same coordinates (batch), each pair (images from same device) or each image.
    (2) Images are loaded and a crop window is drawn and selected. This crop is applied to all related images, depending on selection. Images are saved as TIF files in the “crop” folder.
    (3) An Optional checkbox to use crop images for downstream analysis. Selecting will change the main directory to the “crop” folder.
  c. **Segment Tumor Images** This module segments tumor portions from images using an iterative minimum cross-entropy thresholding algorithm Li and Lee (1993). Steps:
    (1) Images are thresholded and converted to masks.
    (2) Segmented images are saved in a “Segmented” sub-directory as JPEG files.
    (3) Measurement results (e.g., total area, mean gray value) are saved to a CSV file.
  d. **Segment Tumor Weka** This module segments tumor images using a user-trained classifier model with the Trainable Weka Segmentation tool (Arganda-Carreras et al. (2017)). Steps:
    (1) Images are segmented and converted to binary. They are saved as TIF files in the “Tumor Segmented” folder.
    (2) An optional dialog to use the segmented images in downstream steps is provided, though often not used for tumor images.
  e. **Measure Tumor Gray Level** This tool calculates mean, modal, minimum, and standard deviation of gray values for tumor images. Results are saved in a “measured_gray” CSV file.
  f. **Threshold Vessel Images** Thresholds vessel images using the same cross-entropy algorithm. Steps:
    (1) Images are thresholded and saved as JPEG files in a “Threshold” sub-directory with filenames appended with “_threshold.”
  g. **Segment Vessel Weka** This module segments vessel images from a user-trained classifier model with the Trainable Weka Segmentation tool. Steps:
    (1) Images are segmented and converted to binary. they are saved as TIF files in the “Vessel Segmented” folder.
  h. **Measure Vessel Diameter** This cleans and filters vessel images before running the AnalyzeSkeleton plugin to skeletonized images before quantifying diameter, branch point, number of segments, area, and perimeter.
    (1) Hole Filling Threshold: Default = 50; specifies the maximum size of holes to fill in the image.
    (2) Vessel Area Threshold: Default = 10; defines the minimum vessel area to retain.
    (3) Branch Mean Threshold: Default = 50; sets the minimum mean intensity for branches to be kept.
    (4) Junction Distance Threshold: Default = 10; determines the maximum distance at which junction points are considered duplicates.
    (5) Image Cleaning Threshold: Default = 3; sets the Euclidean Distance Map (EDM) threshold for edge pruning to clean the image.
  i. **Trace and export as.DXF** This tool converts the outlined vessels image into a.DXF file. It follows a similar methodology as Kwiatek (2025). Steps:
    (1) Enable Smoothing. This runs the Shape Smoothing plugin to smooth the contours of binary images.
    (2) This sets the Relative proportion FD percent.
  j. **Perfusion Coefficient** This tool measures extravascular leak. Steps:
    (1) Choose an ROI radius (default: 25 pixels) and specify the number of images in the time course (default: 3).
    (2) Optionally align images manually.
    (3) Place ROIs and confirm placement.
    (4) Results are output to a CSV file, and labeled images are saved in a “Perfusion” folder.
  k. **Perfusion Quantification** This tool measures extravascular leak using a less accurate method more automated process. Steps:
    (1) Set the number of images in the time series.
    (2) Selected the image as reference.
    (3) Results are output to a CSV file.

**Figure 1.**
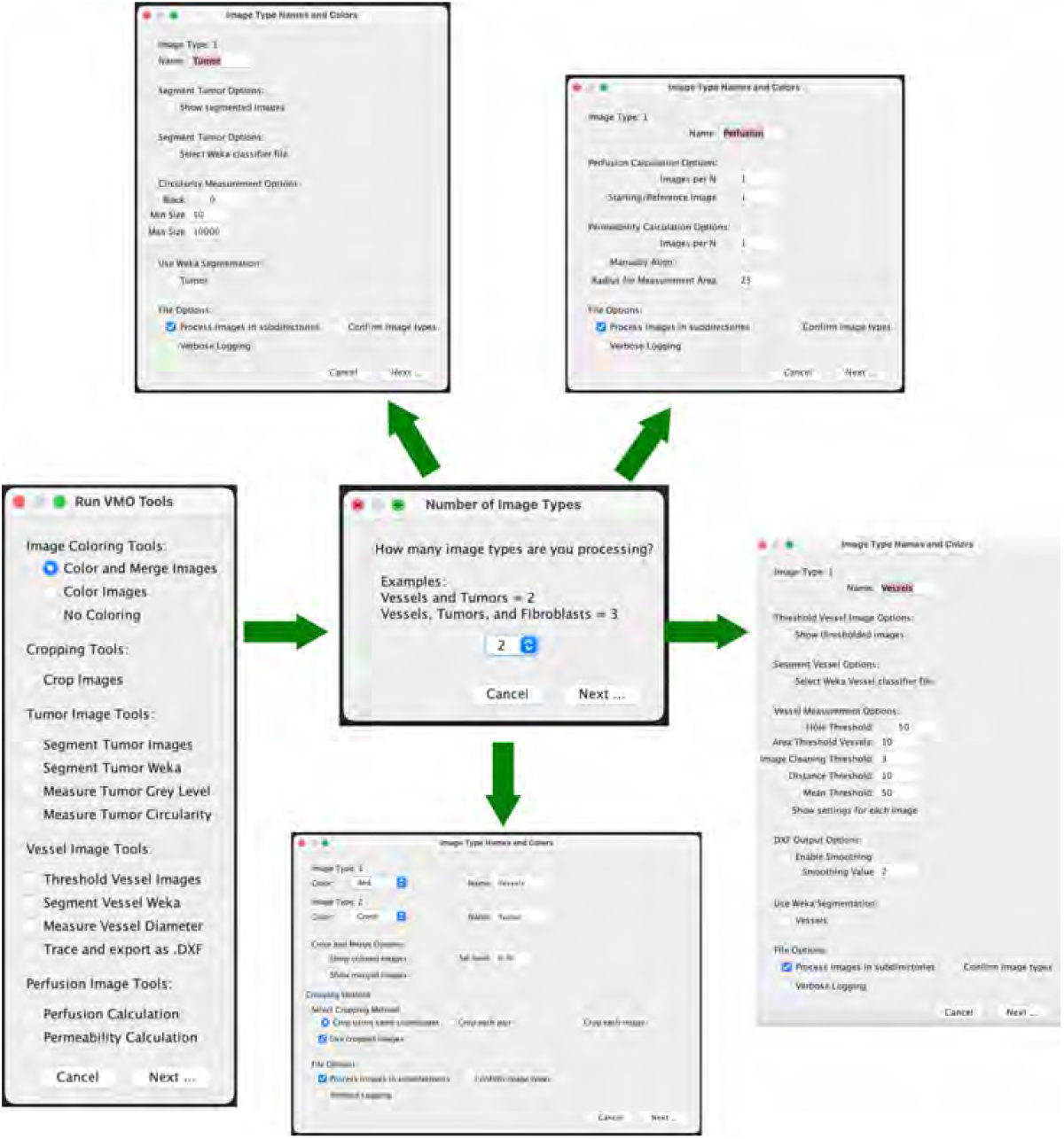
Graphical user interface for Hughes Lab Tools suite. Workflow with options shown.

## 4 RESULTS

### 4.1 Vascularized Micro-Organs and Tumors: Physiologically Relevant Preclinical Models

The VMO/VMT models integrate a living, perfused vascular network that transports oxygen and nutrients to a miniaturized tissue or organ construct. These models have been previously validated as robust in vitro systems for healthy tissue modeling, disease studies, and drug screening applications Sobrino et al. (2016); Phan et al. (2017b); Hachey et al. (2021, 2022, 2023, 2024)). Each high-throughput microfluidic platform incorporates multiple tissue units within a bottomless 96-well plate, enabling independent treatment of each VMO or VMT (Figures 2a, 2b). The device is fabricated from transparent, biocompatible polydimethylsiloxane (PDMS), providing an optically clear platform optimized for real-time microscopic imaging. With each tissue chamber measuring 1 mm^3^ in volume (200 μ*m* deep), only a small number of cells are required for establishment, and minimal reagent volumes are needed for maintenance and therapeutic testing. This setup also facilitates high-resolution imaging using confocal and fluorescence microscopy.

**Figure 2.**
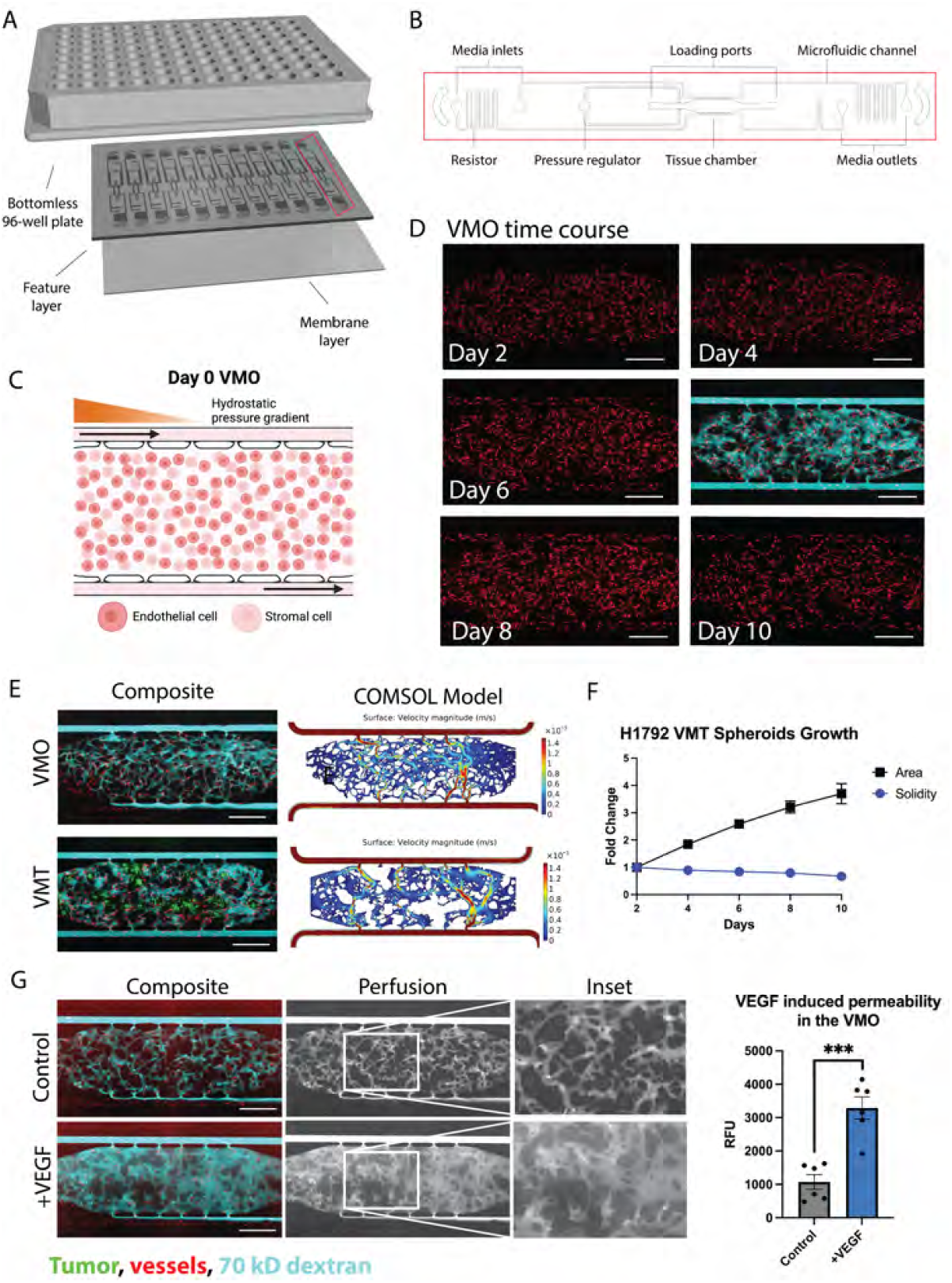
The VMO and VMT as a high-throughput platform for realistic tissue modeling and direct visualization of the vascular niche. (a) Schematic of a microfluidic platform consisting of a bottomless 96–well plate bonded to a feature layer and membrane layer. (b) Schematic of a single device unit with a single tissue chamber fed through microfluidic channels, 2 loading ports (L1-2), and uncoupled medium inlet and outlets (M1-2 and M3-4). A pressure regulator (PR) serves as a burst valve to release excess pressure from the tissue chamber during loading. (c) Schematic showing a zoom view of the chamber loaded on day 0 with endothelial cells and stromal cells, with hydrostatic pressure gradient predominantly from left to right driven across the microfluidic channels. (d) VMO time course of development from day 2 to day 10 with perfusion at day 6. Scale bar = 500 μm. (e) Composite micrographs of VMO and VMT with associated COMSOL models. Tumor shown in green, vessels in red, and 70 kD dextran in cyan. (f) Plot showing H1792 VMT spheroids growth with respect to area and solidity measures. (g) Left: Composite micrographs of perfused VMO (control and VEGF-treated), perfusion only greyscale and inset. Right: Quantification of permeability. *** **p** *<* 0.001

Physiological flow, driven by a hydrostatic pressure gradient across the tissue, enables endothelial cells, stromal cells, and—in the case of the VMT—cancer cells to self-organize within an extracellular matrix, forming a complex microecosystem within five days of culture (Figures 2c, 2d). The resulting vascularized tissue closely mimics an in vivo capillary bed, allowing for physiological drug delivery. To assess vessel patency and permeability changes, vascular networks are routinely perfused with 70 kD FITC- or rhodamine-dextran (Figure 2d). Multiphysics simulations using COMSOL on fully formed, anastomosed, and perfused vascular networks reveal heterogeneous surface velocities of medium flow, closely resembling the dynamic blood flow observed in capillary networks *in vivo* (Figure 2e). In the VMT model, tumor spheroids rely on the microvascular network for nutrient delivery, with their growth and survival closely tied to vascular perfusion. As the spheroids expand within the tissue chamber, they gradually disperse and migrate, leading to an increase in area and a corresponding decrease in solidity, a measure of sphericity, over time (Figure 2f).

Time-lapse imaging of dextran perfusion throughout the tissue chamber allows for the identification of disease-related vascular changes, such as increased permeability or “leaky” vessels in high-grade tumors. To evaluate the responsiveness of microvessels formed within the device, vascular endothelial growth factor (VEGF_165_), a known vascular permeability factor upregulated in the tumor microenvironment and other disease states, was perfused through VMO-associated vessels at 100 ng/mL for 24 hours. Perfusion was then measured using the Perfusion Coefficient tool. Figure 2g illustrates VMO perfusion at 20 minutes under both control and VEGF-treated conditions, with selected region of interests (ROIs) and quantitative analysis confirming the VEGF-induced increase in vascular permeability.

### 4.2 Validation of Hughes Lab Tools for Tumor Measurement: Assessing Tumor Response Across Models

We established VMTs using the H1792 non-small cell lung cancer cell line and evaluated the robustness of the Hughes Lab Tools tumor measurement suite in detecting changes in tumor growth and morphology in response to paclitaxel, a microtubule-stabilizing drug commonly used in advanced non-small cell lung cancer treatment. First, the IC50 in standard 2D monoculture was determined to be 17 nM (Figure 3a). Dose-response experiments in H1792 spheroid monocultures embedded in fibrin showed that paclitaxel treatment (100 nM, 500 nM, and 1 μM) significantly affected both spheroid area (Figure 3b) and solidity (Figure 3c), a measure of sphericity. Over 96 hours, untreated spheroids exhibited a migratory growth pattern, characterized by increased area and decreased solidity. In contrast, treated spheroids remained tightly clustered, with inhibited migration (Figure 3d). Segmentation analysis using the Hughes Lab tool confirmed these distinct growth patterns (Figure 3e).

**Figure 3.**
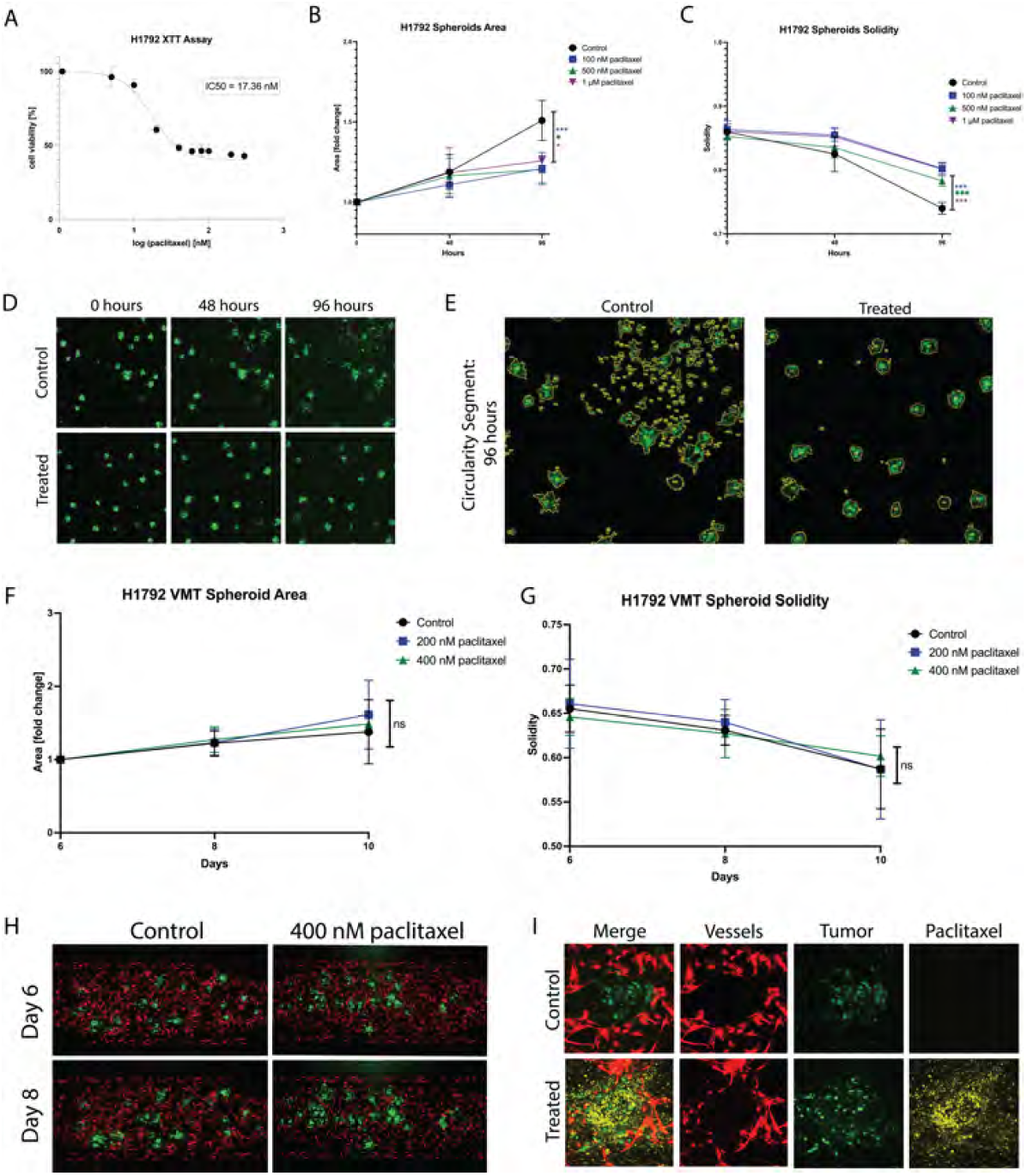
Differential paclitaxel response observed in non-small cell lung cancer spheroids vs. Vascularized Micro-Tumors. (a) Plot showing 2D cytotoxicity results for H1792 treated with paclitaxel for 48 hours. IC50 is 17 nM. (b) Plot showing H1792 spheroid area in response to paclitaxel treatment at 48 hours and 96 hours. (c) Plot showing H1792 spheroid solidity in response to paclitaxel treatment at 48 hours and 96 hours. (d) Micrographs of spheroids (control and 1 μM paclitaxel treated) at time 0, 48, and 96 hours. (e) Segmented tumors at 96 hours for control and treated spheroids. (f) Plot showing H1792 VMT spheroid area in response to paclitaxel treatment for 48 hours at day 6 (baseline), day 8, and day 10. (g) Plot showing H1792 VMT spheroid solidity in response to paclitaxel treatment for 48 hours at day 6 (baseline), day 8, and day 10. (h) Fluorescent micrographs of VMT (control and 400 nM paclitaxel treated) on day 6 and day 8. Tumors shown in green, vessels in red. (i) Confocal micrographs of individual tumor spheroids in the VMT treated with 488 conjugated paclitaxel. Tumor shown in green, vessels in red, and paclitaxel in yellow. ns = non significant, * **p** *<* 0.05, ** **p** *<* 0.01 *** **p** *<* 0.001

In the VMT model, paclitaxel treatment did not significantly affect spheroid area (Figure 3f) or solidity (Figure 3g), as further illustrated by micrographs (Figure 3h). This lack of response was not due to insufficient drug exposure, as VMTs were fully perfused, and confocal microscopy confirmed FITC-conjugated paclitaxel accumulation in the tissue chamber and near tumors within 48 hours post-treatment (Figure 3i). These findings suggest that the complex microenvironment within the VMT may influence drug sensitivity, shifting it toward peak plasma concentrations observed in patients, a phenomenon previously reported by our group (Sobrino et al. (2016); Hachey et al. (2021, 2023, 2024)).

### 4.3 Validation of Hughes Lab Tools for Vascular Morphometry: Comparison with Existing Software and External Datasets

To validate the Hughes Lab Tools vascular morphometry suite, we compared its performance against REAVER (Robust and Efficient Analysis for Vessel Extraction and Reconstruction), a MATLAB-based computational tool for vessel analysis (Corliss et al. (2020)). REAVER has been rigorously tested against other image analysis programs, demonstrating high accuracy and precision across various vessel architecture metrics. Manual measurements served as the gold standard for comparison. As shown in Figure 4a, Hughes Lab Tools achieves vessel segmentation and skeletonization comparable to REAVER when applied to the same raw image file. Error analysis of manual measurements versus outputs from both methods revealed no significant differences in diameter and segment measurements (Figure 4b). Notably, Hughes Lab Tools outperformed REAVER in branch point quantification (Figure 4b).

**Figure 4.**
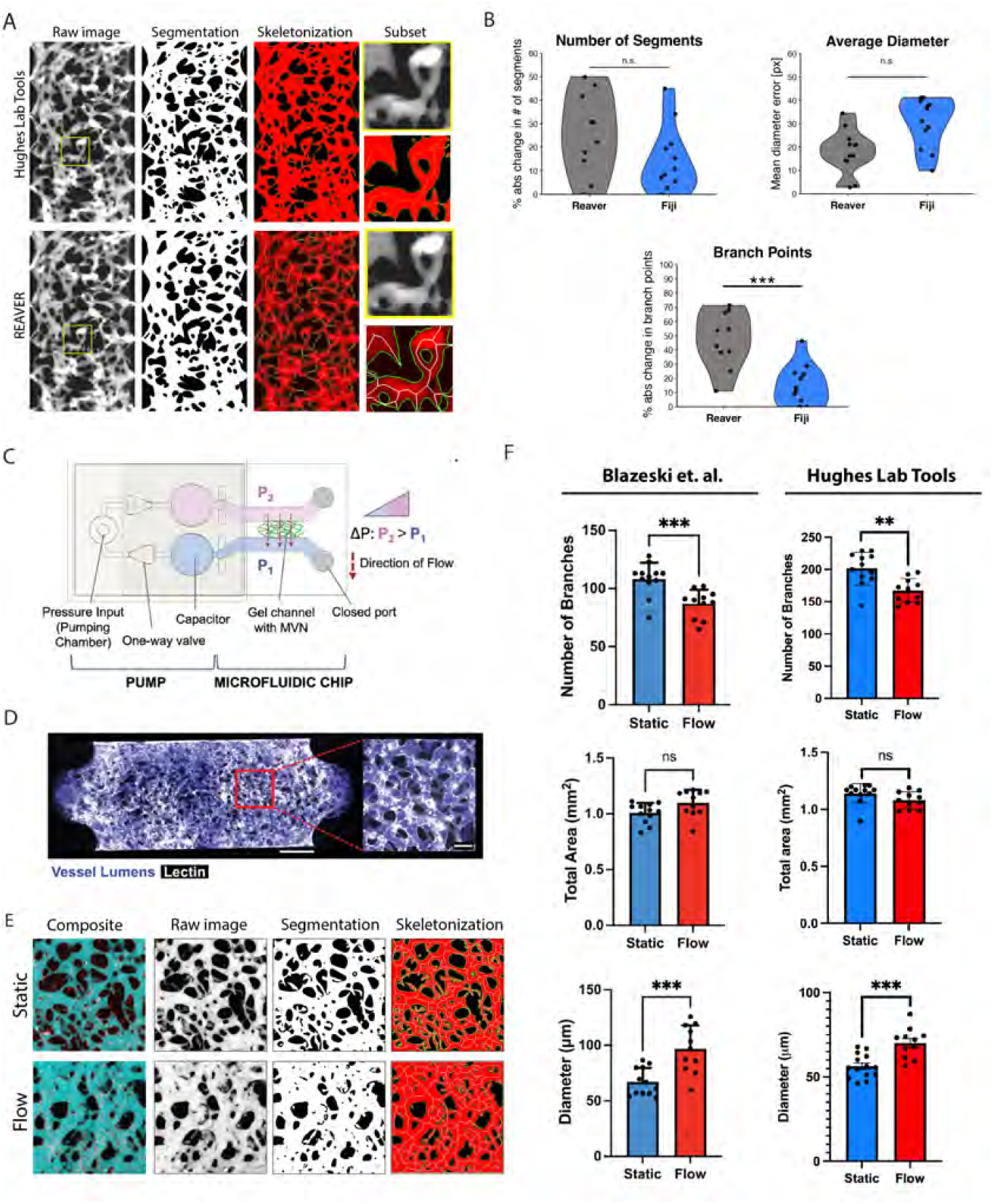
Hughes Lab Tools vascular morphometry suite results. (a) Hughes Lab Tools compared to REAVER for segmenting and skeletonizing raw images of the VMO. (b) Plots showing error comparisons between Hughes Lab Tools or REAVER and manual measurements for branch point, diameter, and segment number measures. (c) Microfluidic chip with pump system for generating microvascular networks (MVNs) (Blazeski et al. (2024)). Reproduced with permission from Elsevier. (d) MVNs perfused with fluorescent dextran (purple) and stained with lectin (white) (Blazeski et al. (2024)). Reproduced with permission from Elsevier. (e) Hughes Lab Tools processing of raw image files from Blazeski et al. (2024), showing composite micrographs of perfused MVNs in either static or flow conditions, segmentation and skeletonization. (f) Plots showing quantification of branch point numbers, vessel diameters, and area between static and flow MVNs, comparing between Blazeski et al. (2024) and Hughes Lab Tools. Reproduced with permission from Elsevier. ns = non significant, * **p** *<* 0.05, ** **p** *<* 0.01 *** **p** *<* 0.001

Further validation was performed using data from an independent research group studying a different organ-on-a-chip model. A recent study by Blazeski et al. utilized a pump-based microfluidic chip (Figure 4c) to generate microvascular networks (MVNs) perfused with fluorescent dextran (Figure 4d) and analyzed using a KLF2-based flow sensor to assess the effects of shear stress on endothelial cell function (Blazeski et al. (2024)). Their findings demonstrated that flow conditions increased vessel diameter, reduced branching, and had no significant effect on total vessel area.

Blazeski et al. analyzed microvascular networks using ImageJ for image segmentation and fluorescent intensity measurements, quantifying total vascular area from maximum intensity projection images of dextran-perfused MVNs. Vessel morphology was assessed with AutoTube, a MatLab-based tool (Montoya-Zegarra et al. (2019)), while the micro-Vasculature Evaluation System algorithm was applied to confocal z-stacks to perform vessel segmentation, skeletonization, and quantify branch number, length, and diameter (Rota et al. (2023)). As shown in Figure 4e, Hughes Lab Tools effectively segments and skeletonizes micrographs from MVNs cultured under static conditions (no flow) and those exposed to flow for 48 hours. Quantitative analysis using Hughes Lab Tools successfully replicates the key findings of the Blazeski et al. study: flow-exposed MVNs exhibit significantly fewer branches than static MVNs, flow conditions lead to a significant increase in average vessel diameter, and total vascular area remains unchanged between the two groups (Figure 4f).

## 5 DISCUSSION

Hughes Lab Tools is an ImageJ suite designed to streamline the processing and analysis of VMO/VMT micrographs, in vitro tumor spheroid models, and other microvascular systems. This tool provides a user-friendly, standardized, and high-throughput solution for extracting imaging data, making it particularly valuable for therapeutic screening and large-scale analyses. Users can specify image locations or process entire subdirectories, enabling rapid analysis of extensive datasets in seconds. The suite also automates image coloring and merging, simplifying data visualization for experimental reference and publication. Benchmark tests highlight its efficiency: on a 2014 MacBook Pro, it processed 600 images (300 tumor/vessel pairs) in just 48 seconds, whereas manual analysis would take over 15 hours, or up to 30 hours with tumor segmentation, demonstrating its transformative impact on imaging workflows.

Beyond automation, Hughes Lab Tools supports longitudinal tumor and vascular analysis, allowing researchers to track changes in spheroid growth, vascular remodeling, and function in VMO/VMT and other preclinical or microphysiological models. A key feature is its vessel morphometry and thresholding function, which prepares images for COMSOL Multiphysics-based fluid flow modeling. The tool processes both total vessel and perfused vessel images, enabling tailored analyses for different experimental objectives, such as evaluating antiangiogenic treatments with total vessels or studying drug or immune cell delivery using perfused vessels. However, differences between thresholded vessel and perfused vessel images may arise from incomplete fluorescent protein transduction in endothelial cells or low-perfusion regions, emphasizing the need for careful experimental design and interpretation.

Validation studies confirm the robustness and broad applicability of Hughes Lab Tools, demonstrating its ability to accurately process external datasets and assess vessel and tumor structures across diverse imaging conditions. Comparative benchmarking revealed that while overall trends were consistent across methodologies, absolute vascular branch counts differed due to sensitivity variations between tools. This discrepancy arises from methodological differences, as Hughes Lab Tools employs REAVER’s model-based approach, which extracts vessel centerlines and estimates radii via intensity profile analysis, making it highly effective for complex vascular networks, including bifurcations and irregular vessel shapes (Corliss et al. (2020)). In contrast, AutoTube assumes a fixed tubular structure, making it better suited for well-defined cylindrical vessels but less precise in complex vascular environments (Corliss et al. (2020)). Our prior work has demonstrated that accurate modeling of small branch points and bifurcations in physiologic vascular networks is crucial for understanding flow dynamics and vascular pruning, underscoring the biological significance of precise vessel segmentation (Hachey et al. (2021); Bender et al. (2024); Hatch et al. (2024)). Importantly, the modular architecture of Hughes Lab Tools allows users to customize functionalities to meet their specific research needs. As an open-source ImageJ-based tool suite, it integrates seamlessly with existing workflows, ensuring accessibility, reproducibility, and efficiency in high-throughput vascular imaging and tumor analysis.

## AUTHOR CONTRIBUTIONS

SJH wrote the manuscript, SJH, CJH, and AGF contributed code, wrote methods, and performed analyses, DG performed experiments, wrote methods, and performed analyses, MLE and JSF contributed code and images, and ALC performed analyses. CCWH provided intellectual input and oversaw the work. All authors contributed to the review and editing of the manuscript.

## FUNDING

This work was supported by the National Institutes of Health, National Cancer Institute and National Center for Advancing Translational Sciences through the following grants: UG3/UH3 TR002137,

R61/R33 HL154307, 1R01CA244571, 1R01 HL149748, U54 CA217378(CCWH) and TL1 TR001415 and W81XWH2110393 (SJH).

## ACKNOWLEDGMENTS

We thank Guillermo Garcia-Cardena (Harvard University) for providing us with the raw image files from his groups’ study (Blazeski et al. (2024)). Figure 2c was created using Biorender.

## REFERENCES

Ahn, S. I., Sei, Y. J., Park, H.-J., Kim, J., Ryu, Y., Choi, J. J., et al. (2020). Microengineered human bloodbrain barrier platform for understanding nanoparticle transport mechanisms. Nature Communications 11, 175. doi:10.1038/s41467-019-13896-7

Arganda-Carreras, I., Kaynig, V., Rueden, C., Eliceiri, K. W., Schindelin, J., Cardona, A., et al. (2017). Trainable Weka Segmentation: a machine learning tool for microscopy pixel classification. Bioinformatics 33, 2424–2426. doi:10.1093/bioinformatics/btx180

Bender, R. H. F., O’Donnell, B. T., Shergill, B., Pham, B. Q., Tahmouresie, S., Sanchez, C. N., et al. (2024). A vascularized 3D model of the human pancreatic islet for ex vivo study of immune cell-islet interaction. Biofabrication 16, 025001. doi:10.1088/1758-5090/ad17d0

Blazeski, A., Floryan, M. A., Zhang, Y., Fajardo Ramírez, O. R., Meibalan, E., Ortiz-Urbina, J., et al. (2024). Engineering microvascular networks using a klf2 reporter to probe flow-dependent endothelial cell function. Biomaterials 311, 122686. doi:10.1016/j.biomaterials.2024.122686

Campisi, M., Shin, Y., Osaki, T., Hajal, C., Chiono, V., and Kamm, R. D. (2018). 3D self-organized microvascular model of the human blood-brain barrier with endothelial cells, pericytes and astrocytes. Biomaterials 180, 117–129. doi:10.1016/j.biomaterials.2018.07.014

[Dataset] Community, G. D. (2025). Git. Accessed: 2025-02-14

Comsol AB (2023). COMSOL Multiphysics® v. 6.3. Stockholm, Sweden. https://www.comsol.com

Corliss, B. A., Doty, R. W., Mathews, C., Yates, P. A., Zhang, T., and Peirce, S. M. (2020). Reaver: A program for improved analysis of highresolution vascular network images. Microcirculation 27, 1–14. doi:10.1111/micc.12618

Ewald, M. L., Chen, Y.-H., Lee, A. P., and Hughes, C. C. (2021). The vascular niche in next generation microphysiological systems. Lab on a Chip 21, 3244–3262. doi:10.1039/D1LC00530H

Gaebler, D., Hachey, S. J., and Hughes, C. C. W. (2024a). Improving tumor microenvironment assessment in chip systems through next-generation technology integration. Frontiers in Bioengineering and Biotechnology 12, 1–27. doi:10.3389/fbioe.2024.1462293

Gaebler, D., Hachey, S. J., and Hughes, C. C. W. (2024b). Microphysiological systems as models for immunologically cold’ tumors. Frontiers in Cell and Developmental Biology 12, 1–21. doi:10.3389/fcell.2024.1389012

Hachey, S. J., Gaebler, D., and Hughes, C. C. W. (2023). Establishing a physiologic human vascularized micro-tumor model for cancer research. Journal of Visualized Experiments, e65865doi:10.3791/65865

Hachey, S. J., Hatch, C. J., Gaebler, D., Mocherla, A., Nee, K., Kessenbrock, K., et al. (2024). Targeting tumorstromal interactions in triple-negative breast cancer using a human vascularized micro-tumor model. Breast Cancer Research 26, 5. doi:10.1186/s13058-023-01760-y

Hachey, S. J. and Hughes, C. C. W. (2018). Applications of tumor chip technology. Lab on a Chip 18, 2893–2912. doi:10.1039/C8LC00330K

Hachey, S. J., Movsesyan, S., Nguyen, Q. H., Burton-Sojo, G., Tankazyan, A., Wu, J., et al. (2021). An in vitro vascularized micro-tumor model of human colorectal cancer recapitulates in vivo responses to standard-of-care therapy. Lab on a Chip 21, 1333–1351. doi:10.1039/D0LC01216E

Hachey, S. J., Sobrino, A., Lee, J. G., Jafari, M. D., Klempner, S. J., Puttock, E. J., et al. (2022). A human vascularized microtumor model of patient-derived colorectal cancer recapitulates clinical disease. Translational Research doi:10.1016/j.trsl.2022.11.011

Hajal, C., Offeddu, G. S., Shin, Y., Zhang, S., Morozova, O., Hickman, D., et al. (2022). Engineered human bloodbrain barrier microfluidic model for vascular permeability analyses. Nature Protocols 17, 95–128. doi:10.1038/s41596-021-00635-w

Hatch, C. J., Piombo, S. D., Fang, J. S., Gach, J. S., Ewald, M. L., Van Trigt, W. K., et al. (2024). SARS-CoV-2 infection of endothelial cells, dependent on flow-induced ACE2 expression, drives hypercytokinemia in a vascularized microphysiological system. Frontiers in Cardiovascular Medicine 11, 1–14. doi:10.3389/fcvm.2024.1360364

Helm, C.-L. E., Fleury, M. E., Zisch, A. H., Boschetti, F., and Swartz, M. A. (2005). Synergy between interstitial flow and VEGF directs capillary morphogenesis in vitro through a gradient amplification mechanism. Proceedings of the National Academy of Sciences 102, 15779–15784. doi:10.1073/pnas.0503681102

[Dataset] Howell, M., Prévost, R., McQuaid, M., and Lalonde, D. (2025). Homebrew. Accessed: 2025-02-14

Hsu, Y.-H., Moya, M. L., Abiri, P., Hughes, C. C., George, S. C., and Lee, A. P. (2013). Full range physiological mass transport control in 3D tissue cultures. Lab Chip 13, 81–89. doi:10.1039/C2LC40787F

Ingber, D. E. (2022). Human organs-on-chips for disease modelling, drug development and personalized medicine. Nature Reviews Genetics 23, 467–491. doi:10.1038/s41576-022-00466-9

[Dataset] Kwiatek, G. (2025). Dxflib, matlab central file exchange. Accessed: 2025-02-26

Li, C. H. and Lee, C. (1993). Minimum cross entropy thresholding. Pattern Recognition 26, 617–625. doi:10.1016/0031-3203(93)90115-D

Lindholm, T., Yellin, F., Bracha, G., and Buckley, A. (2014). The Java Virtual Machine Specification, Java SE 8 Edition (Addison-Wesley Professional), 1st edn.

Low, L. A., Mummery, C., Berridge, B. R., Austin, C. P., and Tagle, D. A. (2021). Organs-on-chips: into the next decade. Nature Reviews Drug Discovery 20, 345–361. doi:10.1038/s41573-020-0079-3

Low, L. A. and Tagle, D. A. (2018). You-on-a-chip’ for precision medicine. Expert Review of Precision Medicine and Drug Development 3, 137–146. doi:10.1080/23808993.2018.1456333

Martier, A., Chen, Z., Schaps, H., Mondrinos, M. J., and Fang, J. S. (2024). Capturing physiological hemodynamic flow and mechanosensitive cell signaling in vessel-on-a-chip platforms. Frontiers in Physiology 15, 1–14. doi:10.3389/fphys.2024.1425618

Montoya-Zegarra, J. A., Russo, E., Runge, P., Jadhav, M., Willrodt, A.-H., Stoma, S., et al. (2019). Autotube: a novel software for the automated morphometric analysis of vascular networks in tissues. Angiogenesis 22, 223–236. doi:10.1007/s10456-018-9652-3

Nahon, D. M., Vila Cuenca, M., van den Hil, F. E., Hu, M., de Korte, T., Frimat, J.-P., et al. (2024). Self-assembling 3D vessel-on-chip model with hiPSC-derived astrocytes. Stem Cell Reports 19, 946–956. doi:10.1016/j.stemcr.2024.05.006

Phan, D. T., Bender, R. H. F., Andrejecsk, J. W., Sobrino, A., Hachey, S. J., George, S. C., et al. (2017a). Blood–brain barrier-on-a-chip: Microphysiological systems that capture the complexity of the blood– central nervous system interface. Experimental Biology and Medicine 242, 1669–1678

Phan, D. T. T., Wang, X., Craver, B. M., Sobrino, A., Zhao, D., Chen, J. C., et al. (2017b). A vascularized and perfused organ-on-a-chip platform for large-scale drug screening applications. Lab Chip 17, 511–520. doi:10.1039/C6LC01422D

Rota, A., Possenti, L., Offeddu, G. S., Senesi, M., Stucchi, A., Venturelli, I., et al. (2023). A threedimensional method for morphological analysis and flow velocity estimation in microvasculature onachip. Bioengineering & Translational Medicine 8, 1–12. doi:10.1002/btm2.10557

Schindelin, J., Arganda-Carreras, I., Frise, E., Kaynig, V., Longair, M., Pietzsch, T., et al. (2012). Fiji: an open-source platform for biological-image analysis. Nature Methods 9, 676

Schneider, C. A., Rasband, W. S., and Eliceiri, K. W. (2012). Nih image to imagej: 25 years of image analysis. Nature Methods 9, 671

Sobrino, A., Phan, D. T. T., Datta, R., Wang, X., Hachey, S. J., Romero-López, M., et al. (2016). 3d microtumors in vitro supported by perfused vascular networks. Scientific Reports 6, 31589. doi:10.1038/srep31589

Urban, G., Bache, K. M., Phan, D., Sobrino, A., Shmakov, A. K., Hachey, S. J., et al. (2018). Deep learning for drug discovery and cancer research: automated analysis of vascularization images. IEEE/ACM Transactions on Computational Biology and Bioinformatics

